# Efficient Sampling of PROTAC-Induced Ternary Complexes

**DOI:** 10.1101/2024.10.30.619573

**Authors:** Hongtao Zhao, Stefan Schiesser, Christian Tyrchan, Werngard Czechtizky

## Abstract

Proteolysis targeting chimeras (PROTACs) are bifunctional small molecules that recruit an E3 ligase to a target protein, leading to ubiquitin transfer and subsequent proteasomal degradation. The formation of ternary complexes is a crucial step in PROTAC-induced protein degradation, and gaining structural insights is essential for rational PROTAC design. In this study, we present a novel approach for efficiently sampling PROTAC-induced ternary complexes, which has been validated using 40 co-crystallized ternary complex structures. In comparison to protein-protein docking-based integrative approaches, our method achieved an impressive success rate of 97% and 50% retrospectively, measured by C*α*-RMSD to the crystal structure within 10 and 4 Å, respectively, with an average CPU time of 4 hours. Notably, utilizing unbound protein structures, the C*α*-RMSD values between the predicted and experimental structures were consistently within 7 Å across six WDR5-PROTAC-VHL ternary structures. Our open-source software enables the modeling of ternary structures in a single step and holds promise for enhancing PROTAC design efforts.

**TOC:** 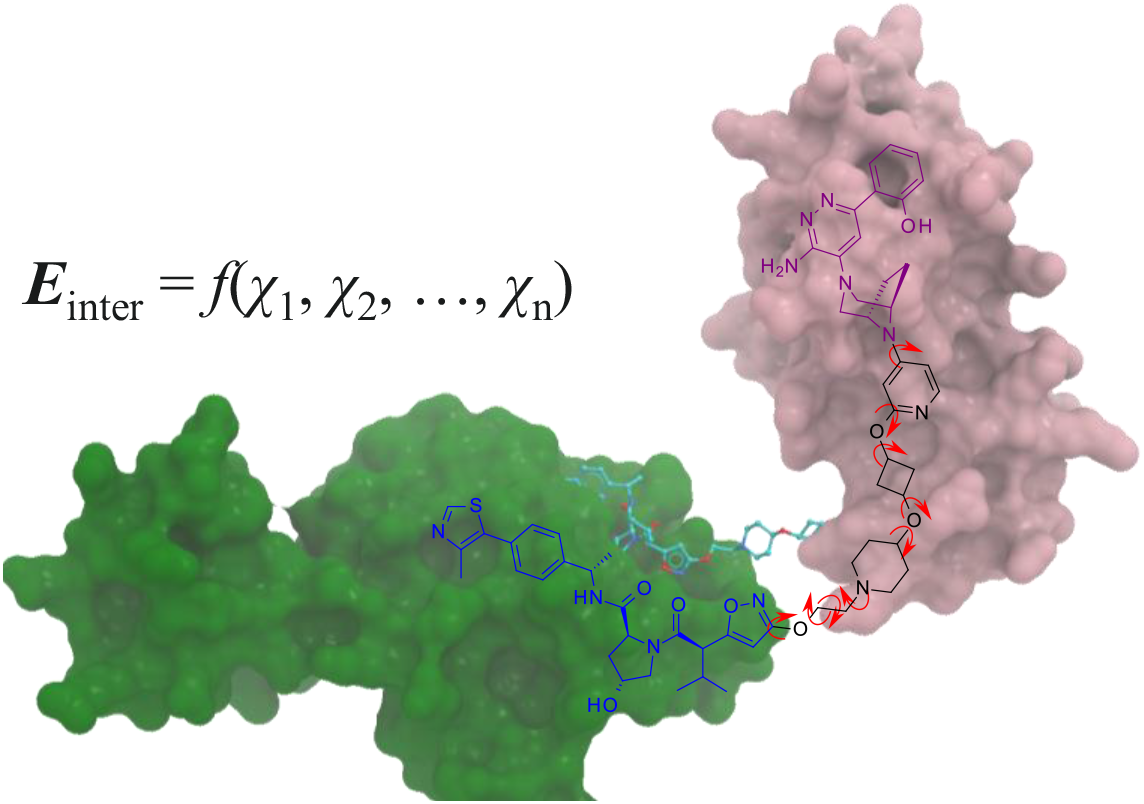

## Introduction

The emergence of targeted protein degradation as a promising therapeutic approach holds the potential to revolutionize the treatment of diseases associated with traditionally undruggable proteins.^1^ Proteolysis-targeting chimeras (PROTACs) represent a critical advance in this field, leveraging heterobifunctional small molecules to recruit specific proteins of interest (POIs) to E3 ubiquitin ligases for ubiquitylation and subsequent proteasomal degradation.^2^ By expanding the druggable proteome, PROTACs offer distinct advantages over traditional inhibition-based therapies.^2, 3^ This innovative approach has rapidly gained momentum, with over 20 PROTACs currently undergoing clinical trials, signaling a paradigm shift in small molecule drug discovery.^4^

Though PROTACs offer conceptual promise, their design poses significant technical challenges, often necessitating the synthesis of numerous analogs to thoroughly explore variations on the linker and exit vectors.^5, 6^ Without extensive trial and error, the selection of optimal linkers and exit vectors remains challenging. On the basis of experimentally determined ternary structures, rational design of selective and potent PROTACs has been demonstrated by identifying new exit vectors or linker rigidification.^7-9^ However, to date, experimental data on PROTAC-induced ternary structures remains scarce, due to the technical difficulties in obtaining high-resolution crystal or cryo-EM structures.^10^ The development of computational methods to model ternary structures is thus in urgent need.

The initial seminal work by Drummond has proposed several methods to model PROTAC-induced ternary complexes,^11^ achieving some success in recapitulating the limited number of available crystal structures at the time of their development.^12-16^ These computational protocols typically begin with protein-protein docking, assuming a rigid-body association of the two proteins. This assumption is supported by experimental structures of existing PROTAC-induced ternary complexes,^17^ while, interestingly, the formation of native protein-protein interactions frequently involves substantial movements or disorder-to-order transitions of the individual proteins.^18^ Furthermore, these methods leverage the knowledge of the bound conformations of the two warheads within their respective proteins. The process involves separate steps for protein-protein docking and linker conformational sampling, with the ternary complex being modeled once a match is identified between them. These methods are computationally demanding, requiring significant resources and time. Furthermore, integrative approaches typically model ternary complexes in a multistep fashion, requiring the use of a variety of toolkits, each of which demands specific expert knowledge. Notably, a recent benchmarking study has shown that neither the MOE approach nor PRosettaC was able to predict ternary structures within 10 Å of the crystal poses for 42% and 58% of the test cases, respectively.^19^

Here, we introduce a computationally efficient method for sampling PROTAC-induced ternary complexes with unprecedented accuracy. Our standalone software, which models ternary complexes in a single step, has been developed using open-source libraries. By sharing our code, we anticipate that it will have a significant impact on PROTAC design.

## Results

### Sampling strategy

We propose a simple method for sampling ternary complexes by considering the interaction energies (*E*_inter_) involved in forming a ternary complex as a function of the rotatable dihedral angles (*χ*) in the linker, as indicated by **Eq. 1**.

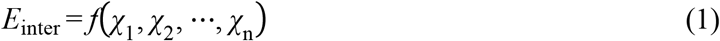

In contrast to previous protocols,^11-16^ this straightforward approach eliminates translational degrees of freedom in protein-protein docking, resulting in a substantial reduction of the search space. The interaction energy can be efficiently evaluated using precomputed grid-based energy maps. Search strategies such as simulated annealing can effectively explore the conformational space to identify energetically favorable ternary complexes. In addition to improving prediction accuracy, our approach enhances computational efficiency by reducing the CPU time from approximately 369 hours for PRosettaC, 53 hours for MOE, and 15 hours for ICM, down to 4 hours (**Table 1**).^19^

**Table 1.** Retrospective Prediction of Near-Native Structures.^*a*^.

| PDB | Resolution | E3 | POI | BSA<sup>b</sup> | $N_{\text{rot}}$<sup>c</sup> | RMSD<sup>d</sup> | $N^e$ | Rank<sup>f</sup> | Time<sup>g</sup> |
| --- | --- | --- | --- | --- | --- | --- | --- | --- | --- |
| 6w7o | 2.17 | BIRC2 | BTK | 1042 | 5 | 5.6 | 5 | 26 | 0.5 |
| 6w8i | 3.80 | BIRC2 | BTK | 194 | 16 | 3.9 | 15 | 17 | 1.1 |
| 8dso | 2.33 | BIRC2 | BTK | 638 | 8 | 8.5 | 14 | 4 | 0.7 |
| 6bn7 | 3.50 | CRBN | BRD4<sup>BD1</sup> | 1049 | 13 | 1.6 | 10 | 18 | 0.6 |
| 6boy | 3.33 | CRBN | BRD4<sup>BD1</sup> | 1133 | 14 | 1.8 | 3 | 22 | 0.7 |
| 6zhc | 1.92 | VHL | BCL-xL | 670 | 20 | 11.1 | 0 | 0 | 1.1 |
| 8fy0 | 2.94 | VHL | BCL-xL | 336 | 9 | 3.3 | 35 | 3 | 0.9 |
| 8fy1 | 2.56 | VHL | BCL-2 | 441 | 9 | 4.5 | 20 | 7 | 0.8 |
| 8fy2 | 2.98 | VHL | BCL-2 | 250 | 6 | 9.1 | 3 | 41 | 0.7 |
| 7pi4 | 2.24 | VHL | PTK2 | 739 | 3 | 4.1 | 504 | 2 | 0.4 |
| 8pc2 | 2.80 | VHL | FKBP5 | 364 | 8 | 1.6 | 40 | 1 | 0.4 |
| 7khh | 2.28 | VHL | BRD4<sup>BD1</sup> | 724 | 12 | 6.6 | 10 | 14 | 0.6 |
| 8bds | 1.72 | VHL | BRD4<sup>BD1</sup> | 624 | 16 | 5.1 | 15 | 2 | 0.7 |
| 8beb | 3.18 | VHL | BRD4<sup>BD1</sup> | 620 | 9 | 3.3 | 34 | 6 | 0.4 |
| 5t35 | 2.70 | VHL | BRD4<sup>BD2</sup> | 730 | 13 | 4.1 | 40 | 4 | 0.5 |
| 7znt | 3.00 | VHL | BRD4<sup>BD2</sup> | 753 | 10 | 0.6 | 22 | 1 | 0.5 |
| 8bdt | 2.70 | VHL | BRD4<sup>BD2</sup> | 724 | 16 | 5.2 | 17 | 6 | 0.8 |
| 8bdx | 2.93 | VHL | BRD4<sup>BD2</sup> | 690 | 16 | 3.7 | 18 | 16 | 0.7 |
| 8r5h | 3.44 | VHL | BRD4<sup>BD2</sup> | 442 | 13 | 6.5 | 20 | 7 | 0.6 |
| 8qu8 | 3.50 | VHL | KRAS<sup>G12V</sup> | 91 | 6 | 4.7 | 5 | 13 | 0.3 |
| 8qvu | 2.24 | VHL | KRAS<sup>G12V</sup> | 494 | 6 | 0.9 | 209 | 2 | 0.3 |
| 8qw6 | 2.20 | VHL | KRAS<sup>G12V</sup> | 609 | 6 | 0.9 | 304 | 1 | 0.3 |
| 8qw7 | 2.36 | VHL | KRAS<sup>G12V</sup> | 566 | 6 | 0.7 | 200 | 1 | 0.3 |
| 6hax | 2.35 | VHL | SMARCA2 | 698 | 6 | 0.5 | 42 | 3 | 0.3 |
| 6hay | 2.24 | VHL | SMARCA2 | 686 | 10 | 4.8 | 11 | 7 | 0.3 |
| 7s4e | 2.25 | VHL | SMARCA2 | 719 | 7 | 8.3 | 3 | 8 | 0.3 |
| 7z6l | 2.24 | VHL | SMARCA2 | 170 | 6 | 2.6 | 29 | 18 | 0.3 |
| 7z76 | 1.32 | VHL | SMARCA2 | 709 | 6 | 0.7 | 81 | 1 | 0.3 |
| 7z77 | 1.97 | VHL | SMARCA2 | 351 | 5 | 2.7 | 57 | 7 | 0.3 |
| 8g1p | 2.70 | VHL | SMARCA2 | 674 | 6 | 2.3 | 31 | 7 | 0.3 |
| 6hr2 | 1.76 | VHL | SMARCA4 | 671 | 6 | 2.9 | 52 | 2 | 0.3 |
| 8g1q | 3.73 | VHL | SMARCA4 | 443 | 2 | 7.3 | 443 | 1 | 0.2 |
| 8qjr | 3.17 | VHL | SMARCA4 | 0 | 9 | 9.2 | 2 | 73 | 0.6 |
| 7jto | 1.70 | VHL | WDR5 | 266 | 15 | 7.7 | 6 | 128 | 1.3 |
| 7jtp | 2.12 | VHL | WDR5 | 831 | 3 | 1.6 | 893 | 1 | 0.4 |
| 7q2j | 2.50 | VHL | WDR5 | 363 | 6 | 1.1 | 220 | 1 | 0.4 |
| 8bb2 | 2.05 | VHL | WDR5 | 100 | 16 | 5.6 | 28 | 14 | 1.0 |
| 8bb3 | 1.80 | VHL | WDR5 | 64 | 16 | 4.2 | 6 | 24 | 1.0 |
| 8bb4 | 2.80 | VHL | WDR5 | 254 | 5 | 2.4 | 27 | 19 | 0.3 |
| 8bb5 | 2.20 | VHL | WDR5 | 181 | 5 | 7.9 | 19 | 4 | 0.4 |

<sup>a</sup>See **Table S1** for linker designations. <sup>b</sup>Buried surface area (Å<sup>2</sup>) of protein-protein interactions. <sup>c</sup>Number of rotatable bonds in the linker. <sup>d</sup>The lowest C $\alpha$ -RMSD (Å) between the POI poses and the crystal structure. <sup>e</sup>Number of POI poses within 10 Å of the crystal structure. <sup>f</sup>Rank of the cluster having at least one POI pose within 10 Å of the crystal structure ordered by the lowest predicted interaction energy in each cluster. <sup>g</sup>Computational time in hours using eight CPU cores.

PROTAC-induced ternary complexes do not necessarily establish favorable protein-protein interactions, a phenomenon known as negative cooperativity.^7, 20^ In one specific case, such as the ternary complex found in the PDB entry 8qjr,^21^ there is a complete absence of protein-protein contact. The presence of negative cooperativity and the lack of protein-protein contacts could potentially constrain the effectiveness of protein-protein docking-based methods, which rely on the assumption of favorable protein-protein contacts driving the formation of ternary complexes.^14^ However, these factors do not pose an issue for our approach. It shall be noted that our approach shares similarities with the implementation in ICM, which has recently been benchmarked against MOE and PRosettaC,^19^ although specific details have not been reported to the best of our knowledge.

### Datasets

The 40 ternary complexes used for validation of this work from PDB (accessed early 2024) correspond to three E3 ligases, namely Von Hippel-Lindau (VHL), Cereblon (CRBN) and Baculoviral IAP repeat-containing protein 2 (BIRC2), each having 35, 2 and 3 structures, respectively (**Figure 1**). The POIs belong to the six protein families, namely bromodomains (BRD4,^7, 20, 22-25^ SMARCA2^8, 26-28^ and SMARCA4^8, 21, 28^), tyrosine kinases (BTK^29, 30^ and PTK2^31^), BCL-2 family proteins (BCL-xL^32, 33^ and BCL-2^33^), immunophilins (FKBP5^34^), small GTPases (KRAS^35^), and WD repeat proteins (WDR5^36, 37^). The resolution of the structures ranges from 1.32 to 3.8 Å, with a median of 2.22 Å (**Table 1**).

**Figure 1.**
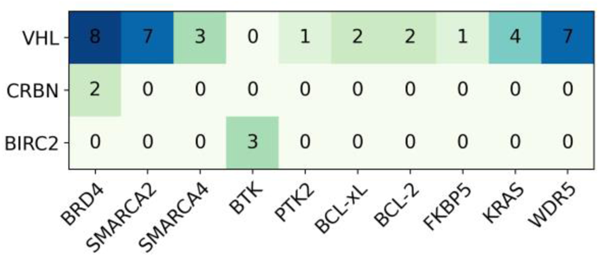
Heatmap depicting the E3-POI pairs in the analyzed crystal ternary complexes.

### Retrospective validation

First, we evaluated performance on the retrospective prediction of near-native ternary structures using bound protein conformations extracted from their respective ternary complexes. Following the previous success metric with C*α*-RMSD to the crystal structure within 10 Å,^11^ our approach achieves a success rate of 97.5%, with 40% predicted at near-atomic resolution around 3 Å (**Table 1**). At a threshold of 4 Å, consistent with accepted criteria in protein-protein docking,^38^ it achieves a 50% success rate (**Figure 1A**).

To put C*α*-RMSD values into context, we looked into crystal ternary complexes featuring the same E3-POI pair induced by the same PROTAC molecule. Among the 40 ternary structures, two pairs of VHL-BRD4^BD2^ induced by **MZ1** and VHL-KRAS induced by **ACBI3** were identified (**Figure 3**).^7, 25, 35^ The two structures exhibit close similarity, with a C*α*-RMSD value of 2 Å between the POIs aligned with the E3 ligase (**Figure 3A**),^7, 25^ while the two proteins share a comparable orientation with a similar protein-protein interface, indicated by a C*α*-RMSD value of 8 Å (**Figure 3B**).^35^ Note that all four structures were experimentally determined. The existence of multiple ternary complexes is attributed to conformational dynamics of PROTAC-induced ternary complexes.^27, 29, 39^

After observing successful performance in recapitulating crystal structures, we focus on understanding how a near-native structure is ranked. A cluster is considered near-native if it contains at least one structure with a C*α*-RMSD within 10 Å to the crystal conformation. Approximately 20% of the near-native clusters were ranked on top, and 45% were within the top five choices, ordered by the predicted interaction energies (**Figure 2B**). The relatively poor ranking of near-native structures could be attributed to the inaccuracy of the underlying scoring function, as well as potentially being influenced by the conformational dynamics of the ternary complex.^27, 29, 39^ The crystal structure could be biased by artifacts arising from crystal contacts, especially for ternary complexes with minimal protein-protein contacts exhibiting the buried surface area (BSA) below 100 Å^2^ (**Table 1**).

**Figure 2.**
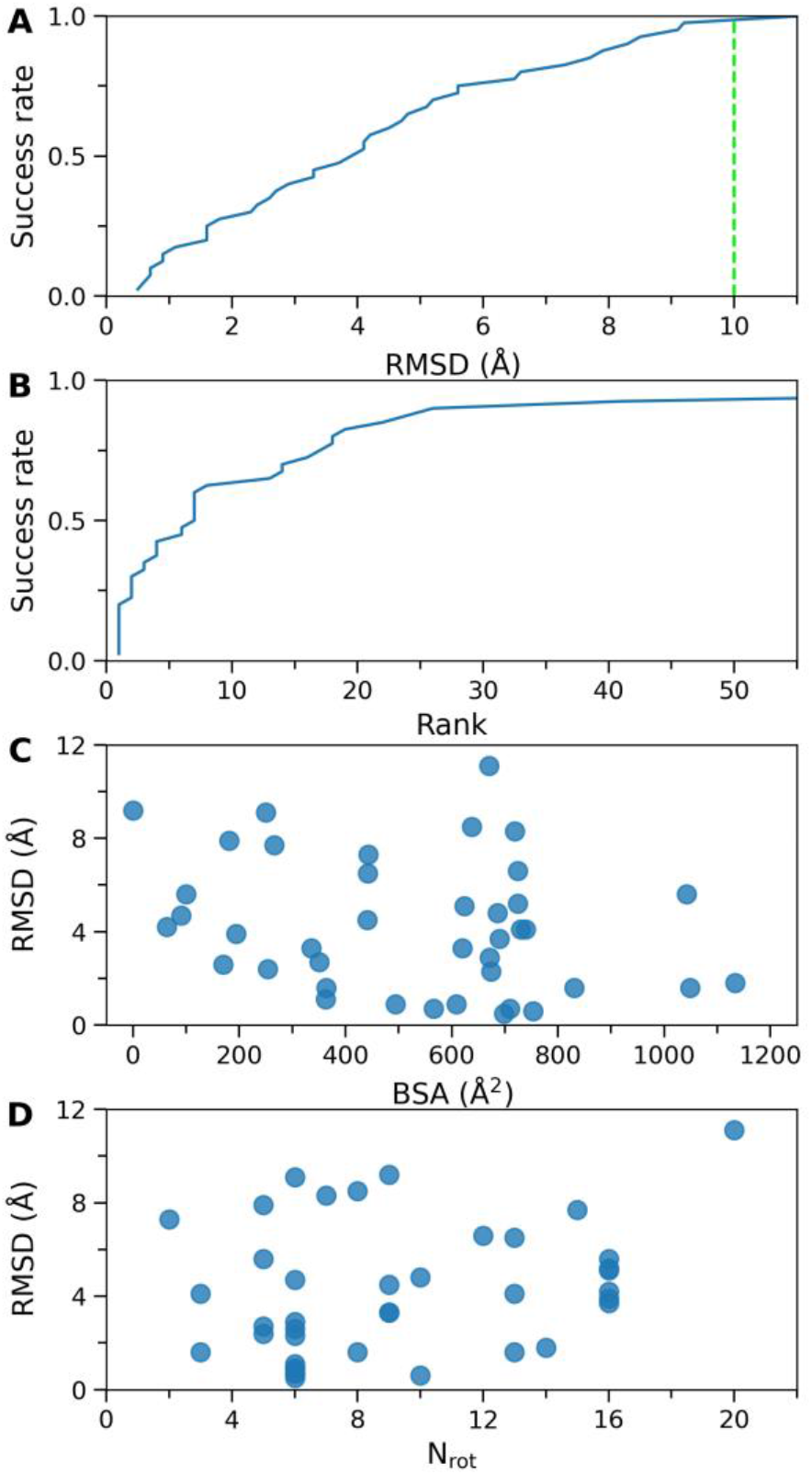
Performance on the retrospective prediction of near-native ternary structures. Cumulative distribution of the lowest C*α*-RMSD (**A**) and best rank of the cluster having at least one near-native pose (**B**), and scatter plots of RMSD against buried surface area (**C**) and number of rotatable bonds in the linker (**D**). The green line corresponds to the cutoff defining a crystal-like pose.

**Figure 3.**
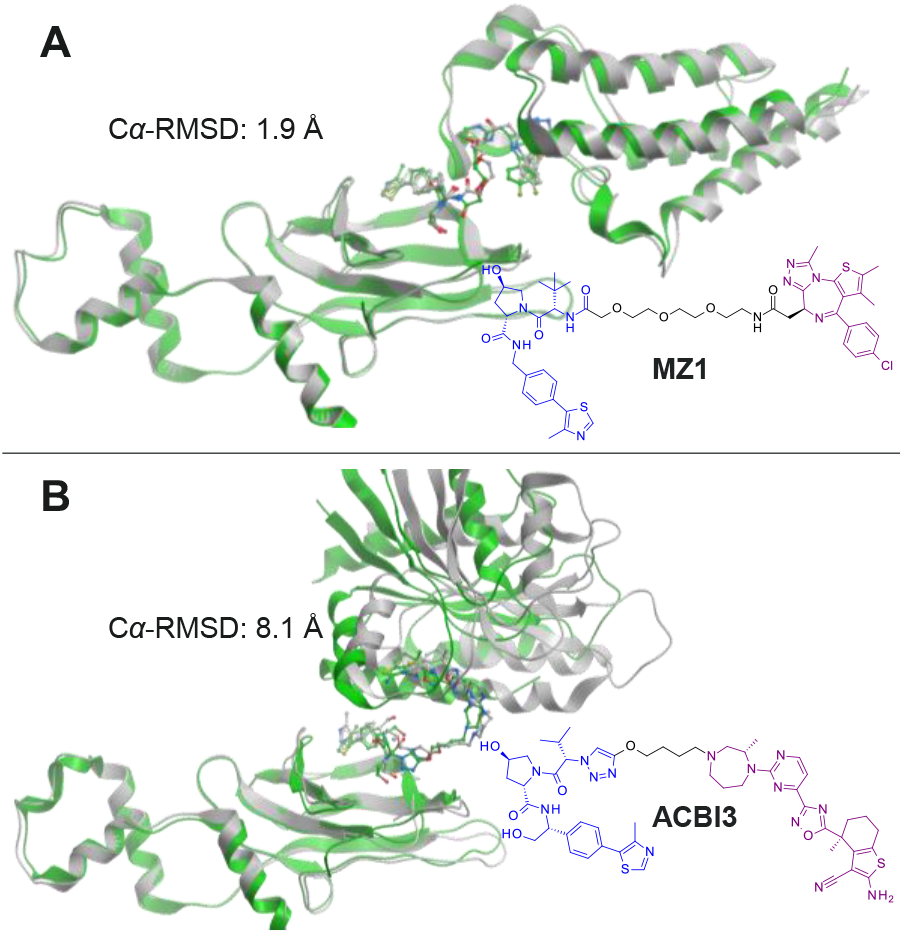
Conformational dynamics of PROTAC-induced ternary complexes. **A**) VHL-**MZ1**-BRD4^BD2^ ternary complexes from the PDB entry 5t35 (colored in green) and 8r5h (in gray) and **B**) VHL-**ACBI3**-KRAS from the entry 8qvu (in green) and 8qu8 (in grey) superimposed on VHL. The C*α*-RMSD values between the POIs are shown.

The BSA resulting from protein-protein interactions varies from 0 to 1133 Å^2^.^17^ Notably, there is no correlation between the lowest C*α*-RMSD values and the BSA, except for the four structures with a BSA below 100 Å (**Figure 2C**). The C*α*-RMSD values for these four structures were above 4 Å, suggesting potentially greater conformational dynamics and a greater influence from crystal contacts. It is noteworthy that the PROTAC in 8qjr degraded SMARCA4 with a DC_50_ of 5 nM, despite the absence of protein-protein contact in the crystal ternary structure.^21^ This raises the question if protein-protein interactions are essential for ubiquitin transfer, or if the observed ternary structure represents a productive state among the ensemble of possible structures. The number of rotatable bonds in the linkers ranges from 2 to 20 with a median of 8. No correlation between performance and the number of rotatable bonds was observed either (**Figure 2D**). In general, an increase in the number of rotatable bonds expands the search space and can make it challenging to converge to the best solution, as may be the case for the structure in 6zhc.^32^ The averaged success rate over three replications is 97%, highlighting the robustness of our sampling approach (**Table S1**).

### Prospective prediction

Next, we aimed to assess how our approach performs in replicating crystal ternary structures, beginning from unbound protein conformations. We have chosen six WDR5-PROTAC-VHL structures, corresponding to five PROTACs that differ only in the exit vector or linkers (**Figure 4**).^40^ This enables us to assess the ability of predicted interaction energies in ranking DC_50_. Encouragingly, the C*α*-RMSD values for all six structures were within 7 Å, and three of them were below the accepted CAPRI criteria of 4 Å in protein-protein docking (**Table 2**).^38^ The computational efficiency of our sampling approach allows for an ensemble of protein conformations to be used in modeling ternary structures. We acknowledge that cross-docking would be a crucial topic that involves varying levels of expert knowledge in the selection of protein conformations. However, the comprehensive evaluation of our approach is beyond the scope of the current study. Leveraging our open-source codes, we aspire for the community to develop enhanced methods for addressing side-chain flexibility at the protein-protein interface, akin to induced-fit docking of small molecules.

**Table 2.** Prospective Prediction on the Six WDR5 Complexes.^*a*^.

| Cpd | pDC<sub>50</sub> | *N*<sub>rot</sub> | PDB | RMSD<sup>b</sup> | *E*<sub>inter</sub><sup>c</sup> |
| --- | --- | --- | --- | --- | --- |
| **1** | 7.3 | 6 | 7q2j | 3.3 | -56 ± 2 |
| **2** | 6.9 | 5 | 8bb4 | 3.5 | -66 ± 0 |
| **3** | 6.6 | 15 | 7jto | 6.0 | -52 ± 4 |
| **4** | <5 | 5 | 8bb5 | 6.8 | -51 ± 1 |
| **5** | <5 | 16 | 8bb2 | 4.9 | -55 ± 2 |
| **5** | <5 | 16 | 8bb3 | 3.9 | -55 ± 2 |

<sup>a</sup>The ternary complex modeling were based on the protein conformation in the PDB entry 4ql1 for WDR5 and 5nvx for VHL. <sup>b</sup>The lowest C $\alpha$ -RMSD (Å) between the POI poses and the crystal structure. <sup>c</sup>The lowest predicted interaction energy in kcal/mol among all poses averaged over three replica runs.

**Figure 4.**
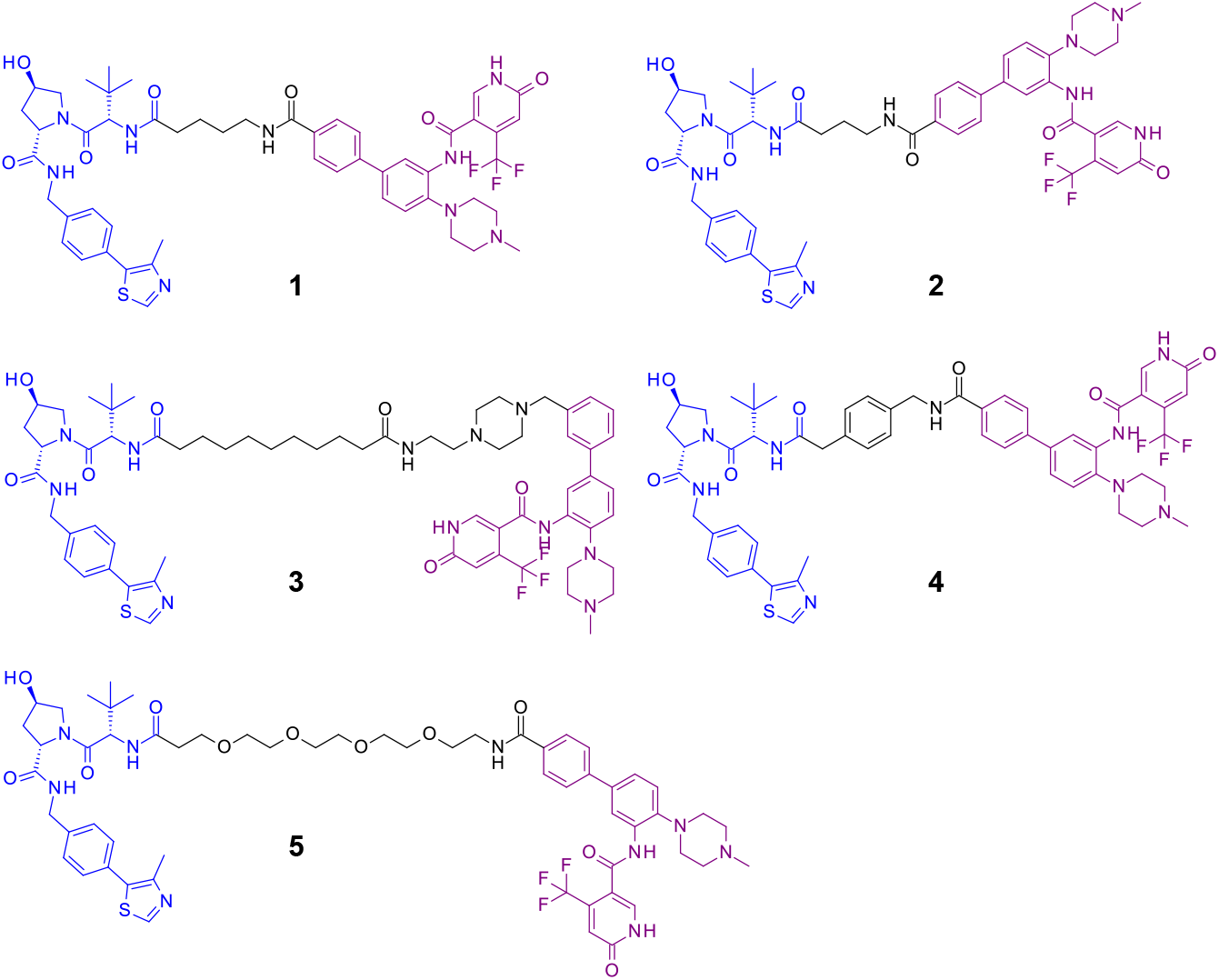
Five WDR5 PROTACs differing in the linker or exit vector. The VHL and POI warheads are colored in blue and purple, respectively.

Maybe not so surprisingly, we did not observe a strong correlation between the predicted interaction energies and DC_50_. It is important to note that DC_50_ is also influenced by the ubiquitylation rate,^41^ which is related to the positions of accessible lysines to the active site of the E2 enzyme, as well as the conformational dynamics of the ternary structures.^17, 27, 29, 39^ Cell permeability is another crucial factor that determines degradation potency.

## Discussion

Growing evidence supports the existence of conformational dynamics in PROTAC-induced ternary complexes, as indicated by both experimental and computational findings.^14, 27, 29, 39^ The crystal structures may be influenced by crystal packing and might not necessarily represent the most functionally relevant complex for ubiquitin transfer. Computational sampling of PROTAC-induced ternary complexes thus offers a valuable approach to comprehend PROTAC-mediated ubiquitin transfer, complementing experimental methods. Additionally, crystal structures have revealed that the same POI could be recruited to the same E3 ligase in diverse orientations, utilizing different surface patches around the E3 binding site. This indicates that observed ternary complexes could be highly specific to individual PROTACs, potentially limiting the utility of crystal structures in the design of novel PROTACs that explore alternative ternary structures. Molecular docking of PROTACs into ternary structures present an additional challenge.^42^ Therefore, our efficient sampling approach could be particularly valuable in future PROTAC design.

## Methods

The interaction energy of forming a ternary complex consists of protein-protein and protein-PROTAC interactions, and PROTAC conformational strain as well.

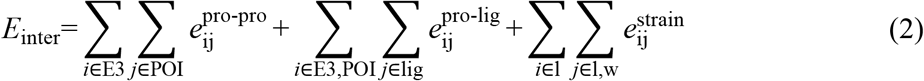

wherein *lig* stands for the PROTAC, *l* and *w* for the linker and the two warheads, respectively. The protein-protein interactions encompass van der Waals, electrostatic, and a linear distance-dependent H-bond energy for donor-acceptor pairs, whereas the protein-PROTAC interactions involve van der Waals and electrostatic interactions only, among heavy atoms. The partial charges of proteins are parameterized using the CHARMM22 force field,^43^ while Gasteiger partial charges are employed for the PROTAC.^44^ In both cases, the partial charges of hydrogens are summed to their attached heavy atoms. The distance-dependent electrostatic energy is further scaled by a coefficient of 0.2. The conformational strain of the PROTAC includes torsion strain and van der Waals interactions within the linker, as well as between the linker and the two warheads, utilizing the Merck force field (MMFF94) with explicit hydrogens.^45^

The conformations of the two warheads in an initial PROTAC conformation are substituted with the bound conformations found within their respective proteins. Upon altering the dihedral angles in the linker, the POI (termed flexible protein) is brought to the proximity of the E3 ligase (termed anchoring protein) by aligning with the bound conformation of its warhead. To speed up the calculation process, grid-based energy maps are precomputed on the E3 for van der Waals, electrostatic and H-bond interactions. For van der Waals interactions, only one atom type (*ε* = 0.1 kcal/mol and *r*_min_ = 1.8Å) is taken into account, utilizing the CHARMM formula.^43^ This enables a rapid calculation of the protein-protein and E3-PROTAC interaction energies. Likewise, grid-based energy maps are generated on the POI by extending 5 Å from its bound warhead. Upon aligning the PROTAC with the bound conformation of the warhead within the POI, its interaction energy with the POI is calculated. The grid space is configured to 0.3 Å. For computational efficiency, the larger protein shall be designated as the anchoring protein. In this study, the E3 ligase was selected as the anchoring protein for benchmarking purposes.

Initially, 10000 linker conformations are randomly generated by uniformly sampling each dihedral angle between ±180°. From these, the 1000 top-scoring conformations are subject to optimization by the simulated annealing approach, starting at a Boltzmann temperature (*k*T) of 60 kcal/mol and subsequently decreasing stepwise by 10 kcal/mol down to 10 kcal/mol. At each temperature, 200 Monte Carlo steps are searched, followed by an energy minimization with the modified Powell algorithm.^46^

The PROTAC conformations in this study were generated using Maestro from Schrodinger, and no modifications were made to the proteins, such as adding missing side chains or loops. This deliberate choice was made to ensure reproducibility by others, as protein preparation can involve varying levels of expert knowledge. However, it is advisable to properly prepare proteins in prospective applications.

## Notes

H.Z., S.S., C.T. and W.C. are employees of AstraZeneca and may own stock or stock options.

## Supporting Information

List of the 40 PDB entries along with DC_50_ values and chemical structures of the corresponding PROTACs, and lowest C*α*-RMSD values from the three replica runs.

## Data availability

The source code and input files for modeling ternary complexes in the manuscript can be found at https://github.com/WIMNZhao/TERNIFY.

