## Supplementary material for "Efficient Sampling of PROTAC-Induced Ternary Complexes": Table S1

Medicinal Chemistry, Research and Early Development, Respiratory and Immunology (R&I), BioPharmaceuticals R&D, AstraZeneca Gothenburg, Sweden

________________________________________

^*^

**Table S1. Retrospective Prediction of Near-Native Structures.**

| PDB | PROTAC*^a^* | pDC_50_ | *N*_rot_ | RMSD*^b^* | | |
| --- | --- | --- | --- | --- | --- | --- |
|  |  |  |  | Run1 | Run2 | Run3 |
| 6w7o | 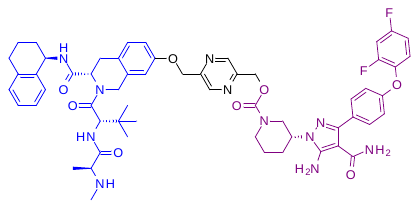 | 6.1 | 5 | 5.6 | 8.1 | 5.8 |
| 6w8i | 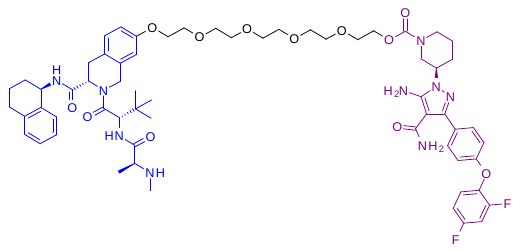 | 6.7 | 16 | 3.9 | 5.4 | 4.5 |
| 8dso | 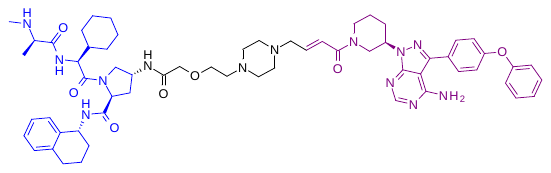 | 7.2 | 8 | 8.5 | 7.5 | 8.3 |
| 6bn7 | 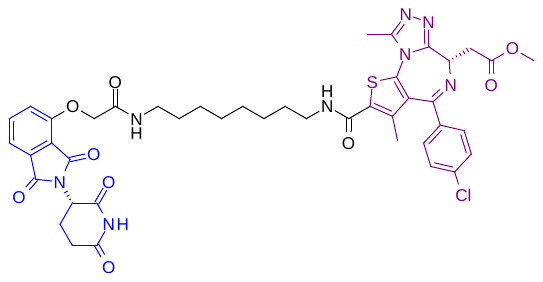 | 7.3 | 13 | 1.6 | 7.1 | 6.6 |
| 6boy | 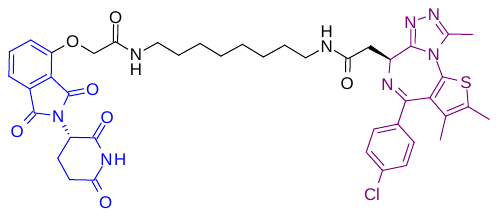 | 8.2 | 14 | 1.8 | 5.7 | 4.0 |
| 6zhc | 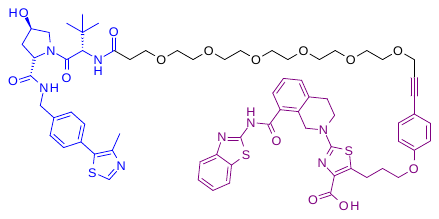 | 8.3 | 20 | 11.1 | 10.8 | 10.3 |
| 8fy0 | 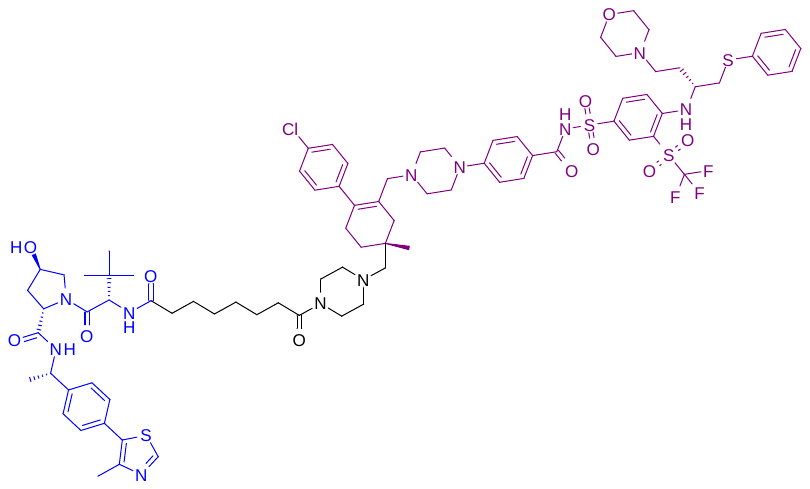 | 8.4 | 9 | 3.3 | 3.4 | 1.6 |
| 8fy1 | 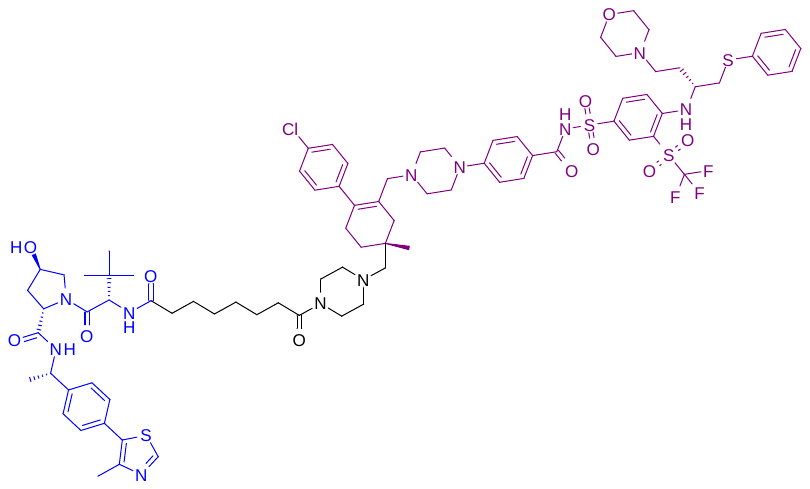 | 7.3 | 9 | 4.5 | 4.7 | 5.8 |
| 8fy2 | 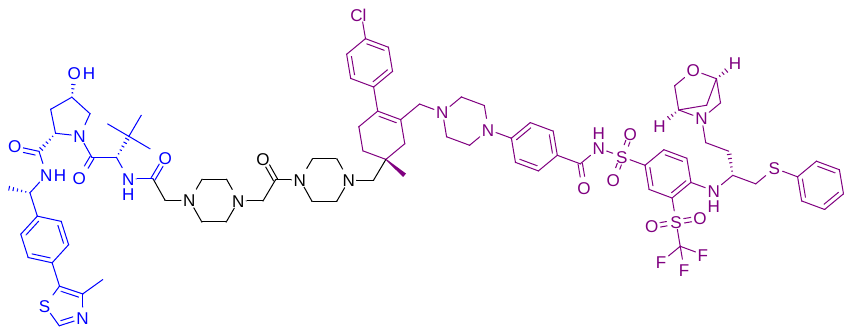 | 8.1 | 6 | 9.1 | 5.2 | 5.9 |
| 7pi4 | 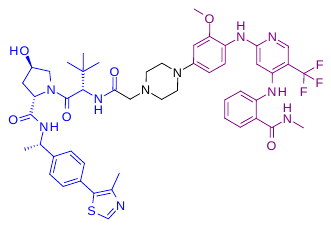 | 8.4 | 3 | 4.1 | 4.1 | 3.9 |
| 8pc2 | 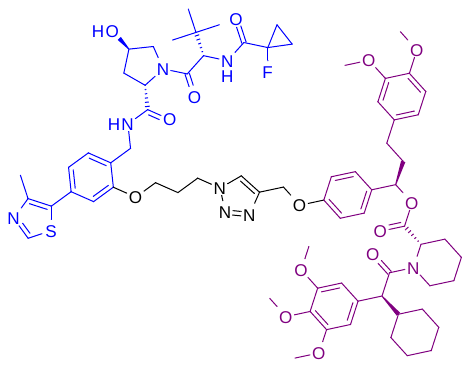 | NA | 8 | 1.6 | 4.1 | 5.6 |
| 7khh | 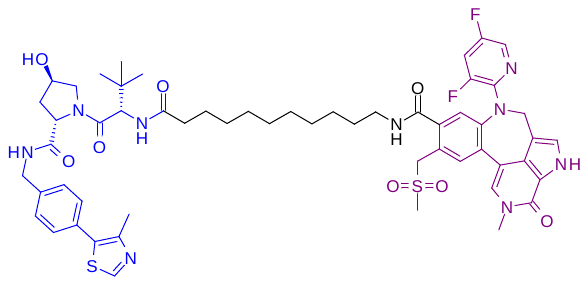 | 10.0 | 12 | 6.6 | 4.3 | 6.0 |
| 8bds | 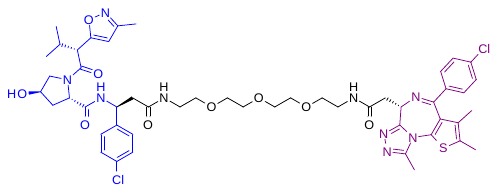 | 6.0 | 16 | 5.1 | 6.4 | 4.1 |
| 8beb | 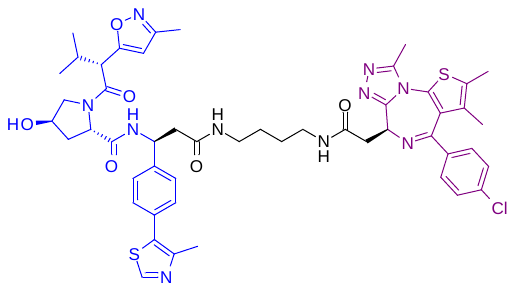 | 6.5 | 9 | 3.3 | 2.0 | 1.1 |
| 5t35 | 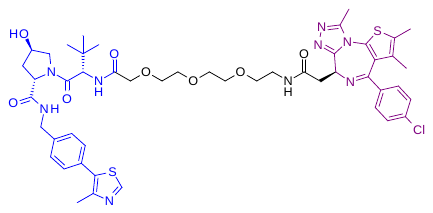 | 8.1 | 13 | 4.1 | 4.7 | 4.6 |
| 7znt | 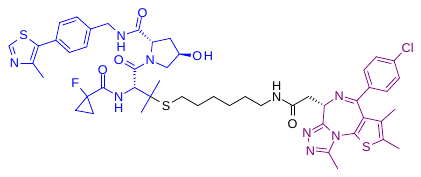 | NA | 10 | 0.6 | 0.8 | 1.0 |
| 8bdt | 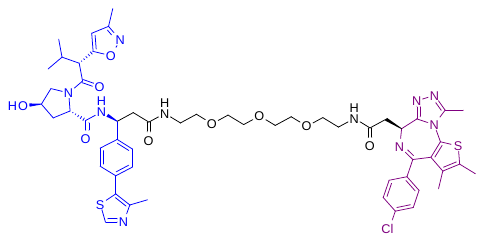 | 6.3 | 16 | 5.2 | 5.9 | 4.5 |
| 8bdx | 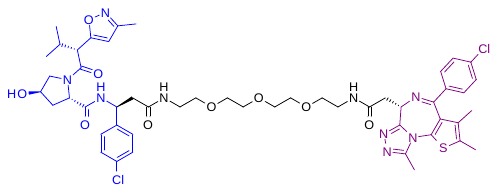 | 6.0 | 16 | 3.7 | 4.0 | 5.5 |
| 8r5h | 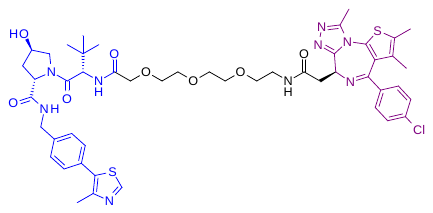 | 8.1 | 13 | 6.5 | 5.5 | 5.5 |
| 8qu8 | 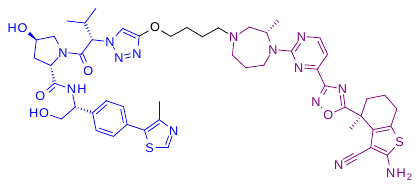 | NA | 6 | 4.7 | 7.5 | 7.5 |
| 8qvu | 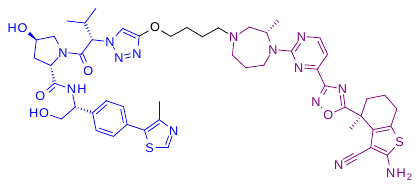 | NA | 6 | 0.9 | 0.8 | 0.6 |
| 8qw6 | 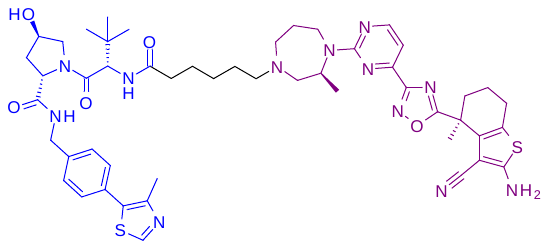 | 6.6 | 6 | 0.9 | 0.7 | 0.8 |
| 8qw7 | 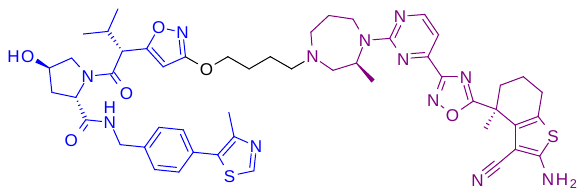 | 7.9 | 6 | 0.7 | 0.6 | 0.7 |
| 6hax | 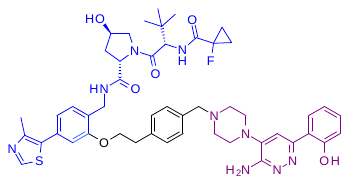 | 7.2 | 6 | 0.5 | 1.1 | 2.6 |
| 6hay | 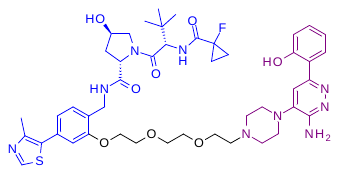 | 6.5 | 10 | 4.8 | 2.3 | 6.0 |
| 7s4e | 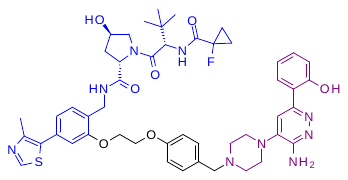 | 8.2 | 7 | 8.3 | 7.1 | 6.9 |
| 7z6l | 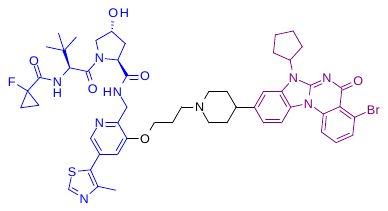 | 7.1 | 6 | 2.6 | 2.1 | 2.1 |
| 7z76 | 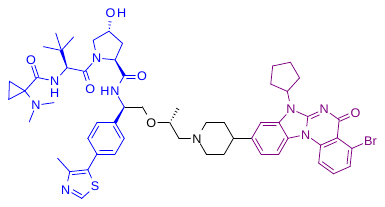 | 8.5 | 6 | 0.7 | 0.4 | 0.8 |
| 7z77 | 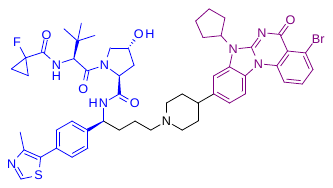 | 8.7 | 5 | 2.7 | 2.6 | 1.9 |
| 8g1p | 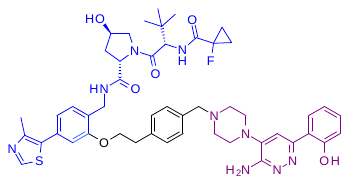 | 6.9 | 6 | 2.3 | 2.4 | 2.7 |
| 6hr2 | 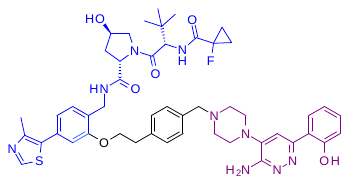 | 6.9 | 6 | 2.9 | 3.5 | 2.2 |
| 8g1q | 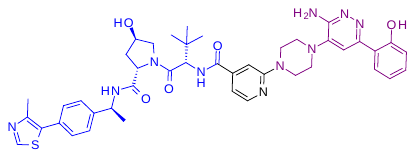 | NA | 2 | 7.3 | 7.3 | 7.3 |
| 8qjr | 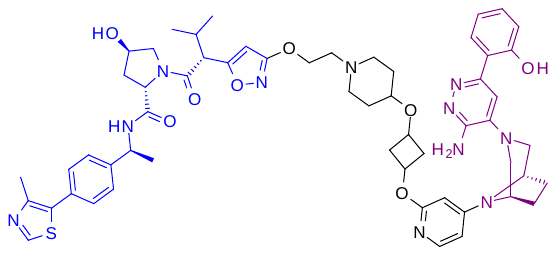 | 8.3 | 9 | 9.2 | 8.4 | 9.1 |
| 7jto | 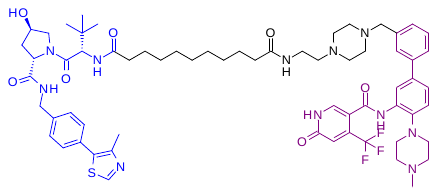 | 6.6 | 15 | 7.7 | 10.2 | 5.3 |
| 7jtp | 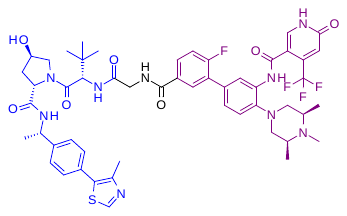 | 8.4 | 3 | 1.6 | 1.7 | 1.6 |
| 7q2j | 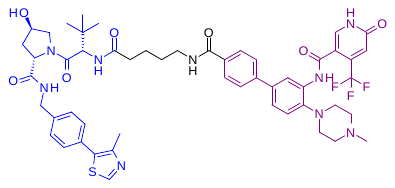 | 7.3 | 6 | 1.1 | 0.7 | 1.0 |
| 8bb2 | 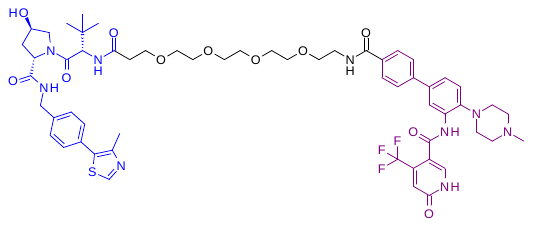 | <5 | 16 | 5.6 | 4.2 | 5.7 |
| 8bb3 | 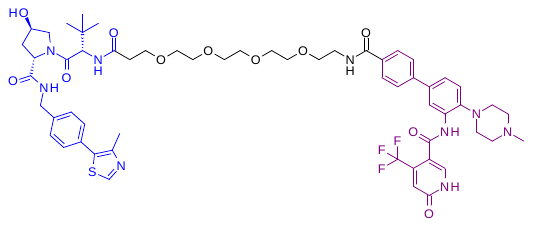 | <5 | 16 | 4.2 | 5.9 | 8.0 |
| 8bb4 | 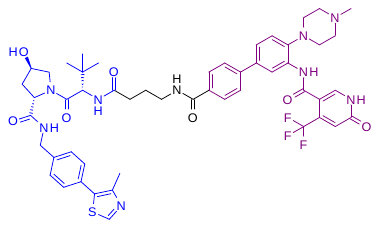 | 6.9 | 5 | 2.4 | 3.4 | 3.2 |
| 8bb5 | 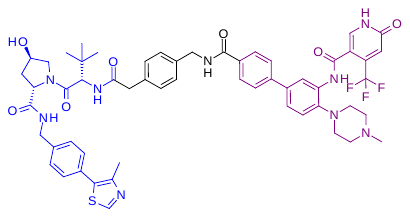 | <5 | 5 | 7.9 | 8.5 | 7.2 |

*^a^*The anchoring (i.e., E3 ligase) warheads are colored in blue and the flexible (i.e., POI) warheads are in purple. *^b^*The lowest C*α*-RMSD (Å) between the POI poses and the crystal structure.
